# An elongated lifespan is associated with higher oxidative protection of flight tissue in Heliconiini butterflies

**DOI:** 10.64898/2026.11.07.765100

**Authors:** Jessica Foley, Pablo Burraco, Stephen H. Montgomery

## Abstract

Evolutionary theories of ageing often invoke trade-offs in resource allocation between somatic maintenance and other important physiological functions. One proposed mechanistic mediator of these trade-offs is oxidative stress, which arises from an imbalance between an organism’s production of reactive oxygen species and its ability to detoxify them. Accordingly, many long-lived species across the animal kingdom show evidence for resistance to oxidative stress. However, it can be difficult to associate lifespan differences with oxidative phenotypes when comparing species across wide taxonomic distances, which may also show large differences in physiology. Butterflies in the long-lived *Heliconius* genus show a 3-fold lifespan extension and slowed ageing in comparison with their close relatives in the Heliconiini tribe, to whom they are phylogenetically, ecologically, and physiologically similar. However, proximate explanations for this extended longevity are currently lacking. Here, we leverage the neat comparative framework between *Heliconius hecale* and *Dryas iulia*, a shorter-lived Heliconiini outgroup, to test whether the observed lifespan differences between the two species are associated with differences in oxidative physiology. Our findings suggest superior oxidative protection of thoracic tissue in the longer-lived *H. hecale* compared to *D. iulia*, as evidenced by increased superoxide dismutase activity, increased total antioxidant capacity, and a corresponding reduction in lipid peroxidation in this tissue. Considering the metabolic demands of insect flight, we suggest that this oxidative phenotype may contribute to the extended longevity of *Heliconius* by enhancing protection of thoracic tissues against oxidative damage, representing a plausible proximate pro-longevity mechanism in this genus.

## Introduction

Understanding why organisms age, and why rates of senescence vary among species, has been a central question in evolutionary biology since the mid-twentieth century (Medawar, 1952; Lemaître *et al*., 2024). Leading theories often place the evolution of senescence within a life history framework (Stearns, 1992), with the disposable soma theory positing that senescence results from trade-offs during the allocation of limited resources between important physiological functions (Kirkwood, 1977). According to this theory, it is the investment of finite resources into reproduction and other physiological processes at the expense of somatic maintenance which ultimately leads to age-related deterioration. Alterations in this resource allocation which allow for increased investment into somatic maintenance may therefore lead to the evolution of lengthened lifespans. This theory has been lent empirical support by the frequent observations of trade-offs between longevity and reproduction in a wide variety of organisms (Harshman and Zera, 2007); however, the physiological mechanisms mediating these evolutionary trade-offs remain poorly understood.

While ultimate explanations for the evolution of ageing were first established many decades ago (Medawar, 1952; Williams, 1957; Kirkwood, 1977; Hamilton, 1966), a more recent preoccupation with the proximate mechanisms of healthy ageing has spurred a growing body of research on unusually long-lived species. Investigations into these species may offer insights into the physiological mechanisms they use to remodel these trade-offs and allow for the evolution of longer lifespans (Rechsteiner *et al*., 2025). One such mechanism thought to play a significant role as a mediator of life-history trade-offs in a wide variety of organisms is oxidative stress (Monaghan, Metcalfe and Torres, 2009). Oxidative stress arises from an organism’s inability to adequately detoxify the excess of reactive oxygen species (ROS) produced as byproducts of mitochondrial respiration, leading to damage of DNA and other biomolecules (Metcalfe and Alonso-Alvarez, 2010). Historically, the age-related increase in ROS production and decline of the capacity to mitigate their damaging effects was thought to be the leading cause of the deleterious effects of ageing (Green, Galluzzi and Kroemer, 2011; Harman, 1956), but a slew of recent studies reporting data contrary to these predictions (Speakman and Selman, 2011) has led to reduced emphasis on oxidative stress as the principal causal mechanism (Speakman *et al*., 2015; Buffenstein *et al*., 2008; Hekimi, Lapointe and Wen, 2011; Lapointe and Hekimi, 2010; Salmon, Richardson and Pérez, 2010). For example, naked mole-rats are born with exceptionally high levels of oxidative damage to their biomolecules (Andziak *et al*., 2006), and yet have lifespans of up to 32 years, ten-fold longer than some other rodent taxa (Edrey *et al*., 2011). One argument settles this apparent contradiction by positing that it is in fact resistance to oxidative stress which confers longevity benefits, as has been suggested in the naked mole-rat (Hulbert, Faulks and Buffenstein, 2006; Edrey *et al*., 2011). In line with this, many other examples of particularly long-lived species show evidence for similarly enhanced stress resistance, including bats (Salmon *et al*., 2009), some long-lived birds (Xia and Møller, 2018; Harper *et al*., 2011; Bize *et al*., 2008), and the clam *Arctica islandica,* the longest-lived non-colonial animal (Ungvari *et al*., 2011). However, the large phylogenetic distances between taxa commonly used in comparative biology of ageing approaches (Gigas *et al*., 2026; Hulbert, Faulks and Buffenstein, 2006; Xia and Møller, 2018) make it difficult to meaningfully interpret observed differences in oxidative physiology. To better elucidate the role of oxidative stress as a potential mediator of life-history constraints, there has been a call for comparisons of oxidative status between closely-related species with differing lifespans (Selman *et al*., 2012).

Butterflies in the long-lived *Heliconius* genus, an emerging model system in the comparative biology of ageing, offer potential empirical evidence for the disposable soma theory. They are among the longest-lived butterflies in the world, and exhibit a dramatic three-fold lifespan extension over their closest relatives in the Heliconiini butterfly tribe (Foley *et al*., 2026). This lifespan extension is associated with the evolution of an adult pollen-feeding behaviour in *Heliconius* which is absent in any other butterfly, including their Heliconiini relatives (Gilbert, 1972). Pollen-feeding provides *Heliconius* with an adult source of amino acids and lipids in addition to the carbohydrate-rich nectar which forms the basis of the diet of the other Heliconiini outgroups. Pollen-derived amino acids are incorporated into *Heliconius* reproductive reserves throughout adulthood (O’Brien, Boggs and Fogel, 2003) and allow for a lengthened reproductive lifespan in *Heliconius* in comparison with their non-pollen-feeding Heliconiini relatives (Dunlap-Pianka, Boggs and Gilbert, 1977). This evolution of an enhanced adult diet appears to have altered the allocation of larval resources away from reproduction and towards somatic maintenance and longevity (Boggs, 1981), with *Heliconius* butterflies showing lengthened lifespans and slowed physiological ageing in comparison with their non-pollen-feeding relatives even in the absence of dietary pollen (Foley *et al*., 2026). While previous research has provided a solid theoretical framework for the evolution of extended lifespan in *Heliconius* (Dunlap-Pianka, Boggs and Gilbert, 1977; Foley *et al*., 2026), mechanistic explanations for how these life-history trade-offs are mediated are currently lacking.

Here, we sought to investigate the possibility that *Heliconius* show increased investment in somatic maintenance, mediated through enhanced resistance to oxidative stress relative to their shorter-lived Heliconiini relatives. The role of oxidative stress may be of particular importance in *Heliconius* considering its relevance to flying insects, which have on average three-fold higher resting metabolic rates than non-fliers (Reinhold, 1999), and where metabolic rates in flight may be up to two orders of magnitude higher than those at rest (Bartholomew and Casey, 1978), contributing to the accrual of oxidative stress (Fisher-Wellman and Bloomer, 2009; Margotta, Roberts and Elekonich, 2018; Levin *et al*., 2017; Du *et al*., 2022; Magwere *et al*., 2006; Yan and Sohal, 2000). By comparing the long-lived, pollen-feeding *Heliconius hecale* and the shorter-lived, non-pollen-feeding *Dryas iulia*, which differ in median and maximum lifespan approximately 2.5-fold (Foley *et al*., 2026), we assessed the degree of oxidative stress at several age points across the captive lifespan (Figure 1). We first measured activity levels of three antioxidant enzymes: superoxide dismutase (SOD), glutathione reductase (GR), and glutathione peroxidase (GPX), as well as total antioxidant capacity (TAC) overall. We also assessed lipid peroxidation as a metric of oxidative damage. To test predicted effects on oxidative phenotypes related to diet and metabolic activity (Monaghan, Metcalfe and Torres, 2009), we also measured these across both pollen-fed and pollen-deprived treatments, taking measurements from both the thorax, which largely consists of the flight muscles, and the abdomen, which largely consists of the gut and reproductive tissue (Figure 1). We hypothesised that the evolution of pollen-feeding in *Heliconius* may have allowed for increased investment of resources towards antioxidant defences, and therefore expected to find evidence for increased antioxidant activity and correspondingly reduced oxidative damage in *H. hecale* in comparison to *D. iulia*.

## Materials and Methods

### Butterfly rearing and sample collection

Stock populations of both *Heliconius hecale* and *Dryas iulia* were first established at the Smithsonian Tropical Research Institute (STRI) in Gamboa, Panama, between January and November 2022, from wild-caught butterflies captured within a 2km radius of Gamboa. Stock populations were maintained as described previously (Foley *et al*., 2026). All known-age experimental individuals were reared from eggs collected from these stock populations, with larvae fed *ad libitum* on leaves of their preferred *Passiflora* host-plants (*P. riparia* for *H. hecale* and *P. biflora* for *D. iulia*) until pupation. Adult lifespan is of specific interest because it is where we see the extension in longevity in *Heliconius* butterflies, with larval and pupal developmental periods being conserved across Heliconiini (Hebberecht *et al*., 2022). “Age” as described in this study therefore refers to adult lifespan, measured in days beginning from time of adult eclosion.

In a previous study we showed median insectary lifespans of 63 days (maximum: 119 days) and 47 days (maximum: 106 days) for *H. hecale* under pollen-fed and pollen-deprived diet treatments, respectively. We found no evidence for an effect of dietary pollen in *D. iulia*, the shorter-lived, non-pollen-feeding Heliconiini outgroup, with an overall median insectary lifespan of 28 days (maximum: 50 days) (Foley *et al*., 2026). Despite this, we considered it still possible that dietary pollen could influence *D. iulia* physiology, given the hypothesis that increased sensitivity to amino acid-rich nectar may represent an early precursor to the evolution of pollen-feeding in *Heliconius* (Young and Montgomery, 2020). We therefore designed our sampling regime as a factorial experiment incorporating species, diet, and age, with an effort made to ensure both sexes were represented across cohorts. To achieve this, we assigned experimental individuals to either a pollen-fed or pollen-deprived diet treatment, and sampled age groups selected to span several age points across the insectary lifespan for each of these species. *D. iulia* was sampled at 3 age points: day 0, day 10, and day 30 post-eclosion, with the distinction between day 0 and day 10 also selected to reflect potential differences related to reproductive effort, as Heliconiini butterflies do not reach sexual maturity until several days post-eclosion (Borrero *et al*., 2026). Similarly, *H. hecale* was sampled at day 0, day 10, day 30, and day 75, designed to mirror the age points in *D. iulia* but adding an additional age point to reflect the longer lifespan in this species, (Foley *et al*., 2026) (Figure 1).

**Figure 1:**
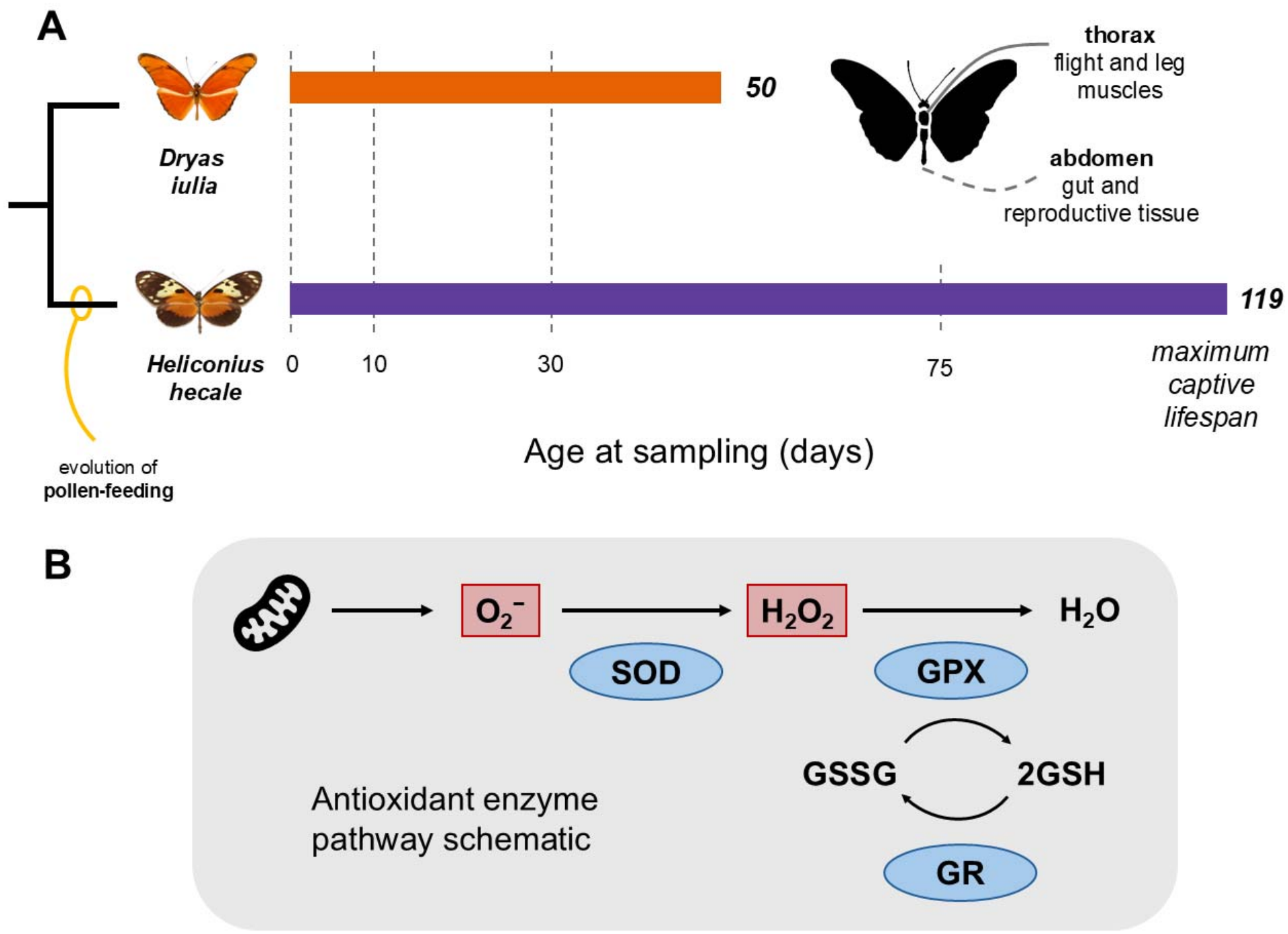
Study design and antioxidant pathway. **A** Species and sampling ages. *Dryas iulia* (orange) and *Heliconius hecale* (purple) are shown on a simplified phylogeny, with bars showing the maximum captive lifespan of each species as recorded in a study conducted under very similar conditions (Foley *et al*., 2026). **B** Schematic of the antioxidant enzyme pathway measured, adapted from (Liu *et al*., 2023). Superoxide radicals (O_2_ ^−^), produced as a byproduct of respiration, are converted to hydrogen peroxide (H_2_O_2_) by superoxide dismutase (SOD). H_2_O_2_ is then reduced to water by glutathione peroxidase (GPX), which oxidises reduced glutathione (2GSH) to its disulphide form (GSSG). Glutathione reductase (GR) regenerates 2GSH from GSSG, allowing for a continuation of GPX-mediated detoxification**. Alt text:** Schematic of study design. **A**: *D. iulia* (maximum lifespan 50 days) and *H. hecale* (119 days) sampled at set ages. **B** Antioxidant enzyme pathway schematic showing roles of the three antioxidant enzymes measured.

Upon eclosion as adults, experimental individuals were weighed, sexed, marked with a unique alphanumeric ID, assigned to an age group and diet treatment, and released into the appropriate experimental cage (pollen-fed / pollen-deprived). A coin-toss was used to assign diet treatment and weighted randomisation with a random number generator was used to assign age group, as older age groups required higher initial populations to account for mortality. To ensure optimal conditions for experimental individuals, and minimise damage to wings from impact with cage walls, we constructed large experimental cages, measuring 4□m (L)□x□3□m (W)□x□2□m (H). Space constraints at the insectaries therefore only allowed for one large experimental cage per diet treatment; however, both species, both sexes, and all age groups were housed together in each of the experimental cages, with overlapping sampling periods, preventing confounding of diet with any other predictor of interest. New experimental individuals were introduced in a staggered manner across the study period, which lasted approximately 6 months from the introduction of the first individual to the sampling of the last individual.

The two experimental cages had matched conditions and resources apart from the provision of pollen. The pollen-fed cage contained four central artificial feeders filled with a 20% w/v sucrose and 10% w/v organic, pesticide-free bee pollen solution, changed every two days. It also contained approximately 12 flowering *Lantana*, *Stachytarpheta,* and *Palicourea* plants, as well as fresh *Psiguria* flowers, replaced daily, to serve as natural floral pollen sources. The pollen-deprived cage contained four central artificial feeders as well as approximately 20 additional artificial feeders placed around the cage, in an effort to standardise the number of flowers and feeders across experimental cages. These artificial feeders were filled with a 20% w/v sucrose solution, changed every two days. The pollen-deprived cage also contained approximately 12 non-flowering *Lantana*, *Stachytarpheta,* and *Palicourea* plants to serve as roosting sites. Both experimental cages contained both *P. biflora* and *P. riparia* host-plants to allow females of both species opportunities to lay eggs.

On the date of sampling (predetermined for each individual based on eclosion date and assigned age group), individuals to be sampled were first weighed on a Sartorius Entris balance (to 1mg) and then decapitated. This meant that total body mass was measured twice for all individuals (with the exception of the day 0 age group): once on the date of adult eclosion, and once more on the date of dissection. The wings and legs were then also removed, and the remaining body was snap-frozen in liquid nitrogen before long-term storage in a −80° freezer, in line with best practices for storage before measurement of biomarkers of oxidative stress (Miranda *et al*., 2026). Individuals were all sampled at the same time of day (15:00, +/– 1 hour) to minimise potential effects on diel variation on oxidative stress markers. Occasionally, time constraints meant that individuals were not sampled on the precise day specified (e.g. sampled at day 31 as opposed to day 30), but this is accounted for in our modelling of age as a continuous parameter. Sampling was conducted in Panama in 2022, and tissues were then transported on dry ice to the U.K. (in 2022) and finally Spain (in 2024) for further analysis under permit number PA-01-ARG-138-2022.

In total, data on body mass was collected for 99 individuals of *H. hecale* and 76 individuals of *D. iulia*. However, tissue samples from many of these individuals were retained for other unrelated analyses of age-related traits. Tissue samples specifically earmarked for the oxidative stress analyses presented in this study were collected for 40 individuals of *H. hecale* and 35 individuals of *D. iulia* in total. For a full breakdown of sample size per species, diet treatment, sex, and age group, see Table 1 (body mass) and Table 2 (oxidative stress)

**Table 1.** Breakdown of sample size for each species/diet/age grouping used for analyses of body mass. Note that for the day 0 group, no diet is assigned, as sampling was conducted before the adult butterfly was provided with food.

| **Age (days)** | ***H. hecale*** |  | ***D. iulia*** |  |
|---|---|---|---|---|
|  | **Pollen-fed** | **Pollen-deprived** | **Pollen-fed** | **Pollen-deprived** |
| **0** | 14 |  | 14 |  |
| **10** | 14 | 16 | 14 | 14 |
| **30** | 14 | 14 | 13 | 21 |
| **75** | 15 | 12 | - | - |

**Table 2.** Breakdown of sample size for each species/diet/age grouping used for analyses of biomarkers of oxidative stress. Note that for the day 0 group, no diet is assigned, as sampling was conducted before the adult butterfly was provided with food.

| **Age (days)** | ***H. hecale*** |  | ***D. iulia*** |  |
|---|---|---|---|---|
|  | **Pollen-fed** | **Pollen-deprived** | **Pollen-fed** | **Pollen-deprived** |
| **0** | 6 |  | 6 |  |
| **10** | 6 | 8 | 6 | 6 |
| **30** | 6 | 6 | 6 | 11 |
| **75** | 6 | 2 | - | - |

### Sample preparation and oxidative stress measurements

Oxidative stress measurements were conducted at the Doñana Biological Station (EBD-CSIC) in Sevilla (Spain) in March 2024. Individual thoraces and abdomens were separated and weighed independently. Tissue samples were homogenised at a 1:4 w:v ratio (tissue mass:buffer volume) in 100 mM Tris-HCl (pH 7.8) containing 0.1 mM EDTA, 0.1 nM PMSF and 0.1% Triton X-100, using a bead mill at 30 Hz for 1 min and subsequently centrifuged at 14,000 rpm for 30 min at 4°C. The resulting supernatants were aliquoted into 1.5-ml microtubes and stored at −80°C until biochemical analyses, which were conducted within two weeks of homogenisation.

We quantified the activities of three antioxidant enzymes, superoxide dismutase (SOD), glutathione peroxidase (GPX) and glutathione reductase (GR), using a COBAS INTEGRA 400 autoanalyser following the protocols described in (Miranda *et al*., 2026). Together, these enzymes form a sequential antioxidant defence pathway in which SOD first catalyses the conversion of superoxide radicals to hydrogen peroxide and oxygen, GPX then reduces hydrogen peroxide and lipid peroxides to water using glutathione, and GR then regenerates reduced glutathione from its oxidised form, allowing for a continuation of GPX-mediated detoxification (Figure 1) (Halliwell and Gutteridge, 2015). GR activity was determined from the decrease in absorbance at 340 nm associated with NADPH oxidation during the reduction of oxidised glutathione (GSSG) to reduced glutathione (GSH), following (Goldberg and Spooner, 1983) and (Melissinos *et al*., 1981). SOD activity was quantified following (Woolliams *et al*., 1983), using the xanthine–xanthine oxidase system to generate superoxide radicals and measuring the inhibition of 2-(4-iodophenyl)-3-(4-nitrophenyl)-5-phenyltetrazolium chloride (INT) reduction. GPX activity was determined following (Paglia and Valentine, 1967), using cumene hydroperoxide as the substrate. In this coupled assay, the GSSG produced during the GPX-catalysed oxidation of GSH is reduced back to GSH by GR, with the concomitant oxidation of NADPH monitored as a decrease in absorbance at 340 nm. Total protein concentration was quantified using the COBAS INTEGRA Total Protein Gen.2 (TP2) assay and used to standardise antioxidant enzyme activities, which were expressed per milligram of protein.

We then quantified total antioxidant capacity (TAC) using the Trolox-equivalent antioxidant capacity assay adapted to the COBAS INTEGRA 400, following (Erel, 2004). The assay measures the inhibition of ABTS oxidation by hydrogen peroxide, using Trolox as the antioxidant standard. Absorbance was measured at 595 nm, and results were expressed as μmol L⁻¹ Trolox equivalents. We also assessed levels of lipid peroxidation by measuring levels of malondialdehyde (MDA), a reactive aldehyde generated during the peroxidation of polyunsaturated fatty acids that is widely used as a marker of oxidative damage to lipids. Lipid peroxidation was estimated by measuring thiobarbituric acid-reactive substances (TBARS), expressed as MDA equivalents, following (Miranda *et al*., 2026) and based on the method of (Buege and Aust, 1978). Briefly, 100 μl of each sample was mixed with 500 μl of an HCl-TCA-TBA reagent containing 0.01% butylated hydroxytoluene. Samples were incubated at 100°C for 15 min and subsequently centrifuged at 10,000 rpm for 10 min at 10°C. The absorbance of the resulting supernatant was measured at 535 nm. MDA concentrations were calculated from a calibration curve prepared using malondialdehyde bis(dimethyl acetal) and expressed as nmol MDA equivalents ml⁻¹. Finally, triglyceride concentration was quantified using the COBAS INTEGRA Triglycerides assay (TRIGL; test ID 0-010).

### Statistical analysis

All statistical analyses were conducted using R v4.4.1 (R Core Team, 2023). In all cases where significant effects of named predictors were found, models including only significant predictors were compared against intercept-only null models to confirm they explained significantly more variance than expected by chance. An analysis of variance (ANOVA) was typically run on models to report significance of all predictors of interest, with any non-significant interactions removed to improve power unless they were of specific interest. Diagnostics for all models were conducted using the using the R package *DHARMa* v0.4.7 (Hartig, 2022). Estimated marginal means and pairwise contrasts were obtained using the R package *emmeans* v2.0.1 (Lenth, 2023), and model predictions for visualisation of results were created with the aid of the R package *ggeffects* v2.3.2 (Lüdecke, 2018).

### Body mass analysis

For all mass analyses, models were fit for two single-species datasets including data across the full lifespan of each species, as well as for an inter-specific dataset with data for both species up to day 30, the oldest age group for which there was data for *D. iulia*. To investigate longitudinal changes in total body mass, the data were first filtered to remove individuals in the day 0 age group, for which only one measurement of total body mass was recorded. Linear mixed models were fit using the package *lme4* v1.1-38 (Bates *et al*., 2015), with individual ID included as a random effect. To accurately estimate within-individual longitudinal declines, we used a within-subject centring approach to separate between-individual effects (e.g. longer-lived individuals tending to be heavier) from within-individual effects (e.g. a longitudinal change in body mass over time), as is recommended for longitudinal analyses (Van De Pol and Verhulst, 2006). We achieved this by creating new variables representing a) mean age for each individual across both weight observations (eclosion/death), and b) deviation from mean age for each observation for each individual, and including these both as predictors in the same model (van de Pol and Wright, 2009). With this approach, a significant effect of mean age is indicative of a between-individual effect (i.e. under representation of smaller individuals in later age groups due to lower survival). Having now controlled for this between-individual effect as a separate term in our model, a significant effect of deviation from mean age can then be taken to indicate within-individual longitudinal change (i.e. decreases in individual body size throughout the life course: our effect of interest in these analyses). A third model including only mean age and actual age as predictors (as well as other variables of interest, e.g. diet, sex etc. as detailed below) was then used to test whether the within- and between-individual slopes differ; a significant effect of mean age in this model indicates that these slopes differ (van de Pol and Wright, 2009). Age was log-transformed to account for non-linear declines in body mass, with a constant (1 day) added to allow for log-transformation of measurements taken at day 0. Single-species models included diet, sex, and their two-way interactions with mean log(age) and deviation from mean log(age) as predictors. Interspecific models additionally included interactions between these predictors and species.

Once we had confirmed declines in body mass with age (see Results), we next sought to separate out the effects of the two tissues of interest (thorax vs. abdomen). As tissue-specific mass had only been measured once for each individual upon dissection, we could not use within-subject centring for this analysis; however, we do not expect this to be a major issue, as only the pollen-fed *H. hecale* cohort showed evidence for selective disappearance (see Results). Linear mixed models were fit using the package *glmmTMB* v1.1.14 (Brooks *et al*., 2017) to model tissue-specific mass as a function of age, diet, sex, whether or not the tissue had been frozen (see Supplementary Note 1), and the two-way interactions of these predictors with tissue, as well as eclosion mass to account for baseline differences in body size between individuals. Tissue mass and eclosion mass were log-transformed to account for allometric scaling and proportional variance in mass. Individual ID was once again included as a random effect to account for shared variance between two tissue measurements from the same individual. Tissue was also included as a dispersion parameter because residual variance in tissue mass differed substantially between the abdomen and thorax.

### Oxidative stress analysis

Before analysing data collected on biomarkers of oxidative stress, several spurious values were removed: five values for TAC which were recorded by the autoanalyser as simply “>4000,00”, as well as one negative value for MDA. We first determined whether activity of antioxidant enzymes was correlated, possibly requiring dimensionality reduction in future analyses, using Spearman’s correlation test as the data were not normally distributed. However, as correlations between antioxidant enzyme activity were not consistent and not particularly strong (see Results), we proceeded with separate models for each enzyme individually. We also tested whether MDA levels correlated with levels of triglycerides, as has previously been found in other taxa (Pérez-Rodríguez *et al*., 2015; Miranda *et al*., 2026). This can bias the reliability of MDA levels, which may then serve to reflect absolute lipid content rather than differences in the proportion of peroxidised lipids (Pérez-Rodríguez *et al*., 2015), and would require inclusion of triglyceride levels as a covariate to account for this. However, Pearson’s correlation test showed no significant correlation between MDA and triglyceride levels in our dataset (*r* = −0.10, *t*_128_ = −1.11, *p* = 0.271) and so we proceeded with further analyses without this covariate.

For each metric of oxidative stress (SOD, GPX, GR, TAC, MDA), single-species generalised linear models were first fit for thorax and abdomen data separately, to investigate effects of predictors of interest within each tissue individually. Because day 0 individuals were sampled before the adult butterfly was provided with food, individuals in this age group did not have an assigned “diet”. Therefore, to explore potential effects of diet, a model including age, sex, and diet was first fit to the subset of individuals aged 10 days and older. If diet did not reach significance in this reduced model, diet treatments were pooled, and a model including only age and sex as predictors was fit to the full dataset (including day 0). We found no significant effect of diet on any biomarker of oxidative stress, and therefore statistics presented on the effects of age and sex in this paper all derive from diet-pooled datasets. Visual inspection of the data revealed no apparent trends in oxidative stress biomarkers with collection date, and collection date was therefore not included as a candidate predictor in subsequent analyses. Inter-specific generalised linear models were then fit for each tissue including species and its two-way interaction with age as candidate predictors to explore differences related to species.

To investigate within-individual differences in oxidative stress across tissues, we then fit single-species generalised linear mixed models for each metric, including tissue as a predictor and individual ID as a random effect to account for shared variance across tissues from the same individual. An inter-specific model was then fit including species and its two-way interaction with tissue as candidate predictors to explore differences related to species. Models were initially fit using the package *lme4* (Bates *et al*., 2015); however, where model diagnostics found heterogeneity in residual variance, models were refit using *glmmTMB* (Brooks *et al*., 2017) to include either tissue or sex as dispersion parameters as necessary.

Models were fit with a gamma distribution for all metrics except for TAC, where gamma-distributed models failed diagnostic checks, and so this metric was therefore log-transformed and modelled using linear models instead. Values for GR contained a substantial number of zeroes (21/150), which corresponded to samples with activity below the assay detection limit rather than technical failures, as confirmed with repeat measurements. Zeroes cannot be modelled using a gamma distribution; therefore, a constant equal to half the lowest detected GR concentration in the dataset (0.01) was added to all GR values prior to analysis. For all models reporting significant effects of predictors of interest, potentially influential observations were assessed either using Cook’s distance (for models fit with *lme4* (Bates *et al*., 2015)) or the testOutliers function from the *DHARMa* package (Hartig, 2022) (for models fit with *glmmTMB* (Brooks *et al*., 2017)). Where observations were identified as potentially influential, either based on a Cook’s distance > 3 x (mean Cook’s distance), or a significant result from the testOutliers function, models were then refit excluding the most influential observation as a sensitivity analysis to assess the robustness of significant effects to its exclusion.

Because five metrics were each tested on the same set of predictors within each species, a Benjamini-Hochberg multiple testing correction was applied to reduce the risk of Type I errors. *p*-values for each species were grouped according to the combination of model type (both-tissues model, thorax model, abdomen model) and predictor (age, sex, diet, tissue, species), with each group comprised of the five *p*-values for that predictor across the five oxidative stress metrics, and the correction applied across those five *p*-values.

Additionally, to test for statistical correlations between TAC and MDA levels within tissues, either Pearson’s or Spearman’s correlation test (for normally or non-normally distributed data, respectively) was performed separately for each tissue, within each of the two species.

### Associations between oxidative stress and change in body mass

We finally sought to investigate whether oxidative stress was associated with the observed declines in body mass. However, as previously noted, we could not observe within-individual change in thorax or abdomen mass directly, and therefore we could not directly map the tissue-specific values for various biomarkers of oxidative stress to changes in tissue mass. We reasoned that an individual’s deviation from the tissue mass expected for its age, sex, and eclosion mass could provide an index of relative tissue preservation or depletion. We took the best-supported tissue-specific mass model for each species and calculated the residual (observed – predicted log mass) for each observation. A positive residual indicates that a tissue was heavier than expected (less depleted), and a negative residual that it was lighter than expected (more depleted). For each species, we tested whether these residuals were associated with biomarkers of oxidative stress by fitting linear and generalised linear mixed models for each biomarker with the model specifications described above. Each biomarker was modelled as a function of the residual, tissue, and their two-way interaction. Individual ID was also included as a random effect to account for shared variance across tissues from the same individual. As described above, the Benjamini-Hochberg multiple testing correction was applied to *p*-values associated with the residual term and its interaction with tissue across the five models for each biomarker. As the residuals used as predictors were derived from a previously-estimated model, they carry their own estimation uncertainty, and so these analyses should be considered exploratory rather than definitive.

## Results

### Age-related declines in body mass driven by species-specific tissue depletion

We previously showed that total body mass declines with age in both *H. hecale* and *D. iulia* (Foley *et al*., 2026). We first confirm these findings in this new cohort by modelling total body mass as a function of age, using a within-subject centring approach to decompose the effect of age into a between-individual component (mean age) as well as a within-individual component (deviation from mean age) (van de Pol and Wright, 2009). The results of these analyses largely recapitulated our previous findings, demonstrating age-related within-individual declines in body mass in both *H. hecale* (Figure 2; χ^2^_1_ = 598.33, *p* < 0.001) and *D. iulia* (Figure 2; χ^2^_1_ = 529.67, *p* < 0.001). In both species, these declines were steeper in males (χ^2^_1_ = 18.83, *p* < 0.001), and in *H. hecale*, but not *D. iulia*, this decline was steeper in pollen-deprived individuals (Figure 2; *H. hecale*: χ^2^_1_ = 9.89, *p* = 0.002; *D. iulia*: χ^2^_1_ = 0.61, *p* = 0.436). These results largely match our previous findings from a separate cohort of butterflies maintained under similar conditions (Foley *et al*., 2026), although where we previously found a linear decline in body mass with age, the declines observed here are non-linear, characterised by a steeper initial loss that decelerated with time (captured by modelling age on a log scale, see Methods). Our analyses also revealed that older pollen-fed *H. hecale* individuals were generally heavier, as the between- and within-subject slopes for this group significantly differed (χ^2^_1_ = 8.27, *p* = 0.004), suggesting that selective disappearance of lighter individuals from older age groups masked a within-individual decline in body mass in this group. However, this was not true of pollen-deprived *H. hecale* (t_87.5_ = 0.392, *p* = 0.6958), nor of *D. iulia*, where the between- and within-subject effects were not significantly different (χ^2^_1_ = 8.27, *p* = 0.004).

Having confirmed our previous finding of age-related declines in body mass in both species, we next investigated whether these declines were driven by depletion of specific tissues – namely, the thorax or the abdomen. In *D. iulia*, both tissues showed declines in mass with age (χ^2^_1_ = 30.14, *p* < 0.001), but rates of decline were similar between tissues (*t*_134_ = 0.59, *p* = 0.558), with both the thorax and the abdomen predicted to lose approximately 22.45% of their mass by day 30 (Figure 2). In *H. hecale*, however, while the abdomen was predicted to deplete by approximately 20.75% by day 75 (*t*_178_ = −3.36, *p* = 0.001), this was not true for the thorax, where the slope was not different from 0 (*t*_178_ = −0.74, *p* = 0.462) (Figure 2). Results from the inter-specific model confirmed that the thorax declined more steeply in *D. iulia* than in *H. hecale* (*t*_274_ = −3.64, *p* < 0.001), although the abdomen slopes did not differ between the two species (*t*_274_ = −1.33, *p* = 0.186).

**Figure 2:**
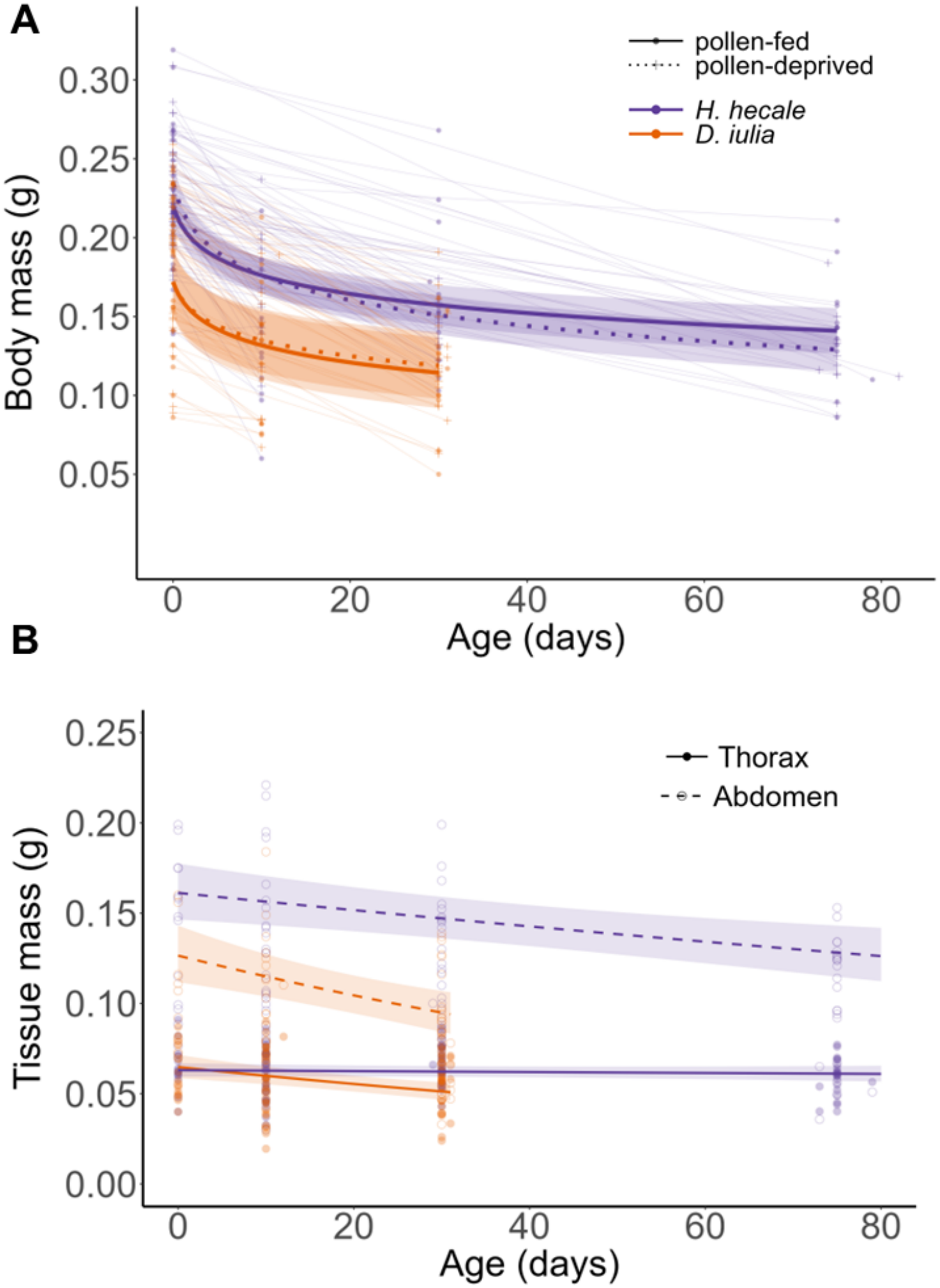
Age-related declines in body and tissue mass. **A** Total body mass (g) of *H. hecale* (purple) and *D. iulia* (orange) across adult life. Points show individual measurements at eclosion and at death, with thin lines joining measurements from the same individual. Thick lines show model-predicted trajectories for pollen-fed (solid) and pollen-deprived (dotted) butterflies, with 95% confidence intervals. **B** Thorax (filled circles, solid lines) and abdomen (open circles, dashed lines) mass (g) at dissection, plotted against age. Lines show predictions from the tissue-mass model with 95% confidence intervals. *n* = 175 individuals (see Table 1). **Alt text**: Line graphs of mass against age. **A**: body mass declines in both species, more steeply early on. **B**: abdomen mass declines in both; thorax mass declines only in *D. iulia*.

### Higher thoracic antioxidant capacity and lower thoracic peroxidation in s

Upon beginning our analysis of oxidative stress biomarkers, we first investigated whether activity levels of the three antioxidant enzymes (GPX, GR, SOD) were correlated, as might be expected considering their involvement in sequential enzymatic reactions (Figure 1). A Spearman’s correlation test showed significant correlations between activity levels of GR and GPX (ρ = 0.36, *p* < 0.001) and GR and SOD (ρ = −0.24, *p* = 0.003), but not between SOD and GPX (ρ = −0.26, *p* = 0.769).

Our analysis of antioxidant enzyme activity and oxidative damage suggested more favourable oxidative status in the longer-lived *H. hecale*. In the thorax, both SOD activity (Figure 3; χ^2^_1_ = 9.16, *p*_adj_ = 0.012) and TAC (Figure 3; *F*_1_ = 6.23, *p*_adj_ = 0.037) were higher in *H. hecale* than in *D. iulia*, with SOD activity increased 1.58-fold and TAC increased 1.36-fold. In accordance with this, *H. hecale* showed evidence for reduced lipid peroxidation in comparison with *D. iulia*, as indicated by a 0.76-fold lower MDA concentration (Figure 3; χ^2^_1_ = 8.52, *p*_adj_ = 0.004). There was no evidence for a significant interaction between species and tissue (χ^2^_1_ = 0.13, *p_adj_* = 0.718), indicating that this reduction in lipid peroxidation in *H. hecale* was consistent across both the thorax and the abdomen. Conversely, levels of GPX activity in the abdomen were 1.59-fold higher in *D. iulia* than in *H. hecale* (Figure 3; χ^2^_1_ = 10.82, *p*_adj_ = 0.005). However, despite this, there was no difference between species in TAC in the abdomen (Figure 3; *F*_1_ = 0.85, *p*_adj_ = 0.451).

Within *H. hecale,* levels of tissue-specific antioxidant enzyme activity did not always correspond; SOD activity was 3.18-fold higher in the thorax than the abdomen (Figure 3; χ^2^_1_ = 118.05, *p*_adj_ < 0.001), whereas GR activity was 3.5-fold higher in the abdomen than in the thorax (Figure 3; χ^2^_1_ = 12.96, *p*_adj_ < 0.001). Despite this, overall TAC was 1.28-fold higher in the thorax than the abdomen in *H. hecale* (Figure 3; χ^2^_1_ = 4.30, *p*_adj_ = 0.047). This pattern is accordingly reflected by the 0.43-fold lower levels of lipid peroxidation in the thorax as compared to the abdomen in *H. hecale* (Figure 3; χ^2^_1_ = 65.87, *p*_adj_ < 0.001), although TAC and MDA levels did not correlate statistically within either tissue (thorax: ρ = 0.27, *p* = 0.149; abdomen: ρ = −0.18, *p* = 0.296). However, this pattern is reversed in *D. iulia*, where despite a nonsignificant trend towards higher TAC in the abdomen compared to the thorax (Figure 3; χ^2^_1_ = 3.02, *p*_adj_ = 0.082), lipid peroxidation was also much lower in the thorax (Figure 3; χ^2^_1_ = 51.11, *p*_adj_ < 0.001), which showed a 0.37-fold lower MDA concentration. In *D. iulia*, TAC and MDA levels also did not correlate statistically within either tissue (thorax: ρ = 0.04, *p* = 0.837; abdomen: *r* = 0.22, *p* = 0.238).

We found no evidence for any meaningful effect of diet or age on antioxidant enzyme activity or oxidative damage in either species. While we did initially find a significant age-related increase in SOD activity in the thorax of *H. hecale* (χ^2^_1_ = 12.07, *p*_adj_ = 0.003), this effect appeared to be driven by a single outlier with high leverage (Figure S1). Removal of this outlier rendered the age effect nonsignificant (χ^2^_1_ = 3.75, *p*_adj_ = 0.264). The only effect of sex was observed in the *H. hecale* abdomen, where males had higher levels of GR activity (χ^2^_1_ = 10.41, *p*_adj_ = 0.006).

**Figure 3:**
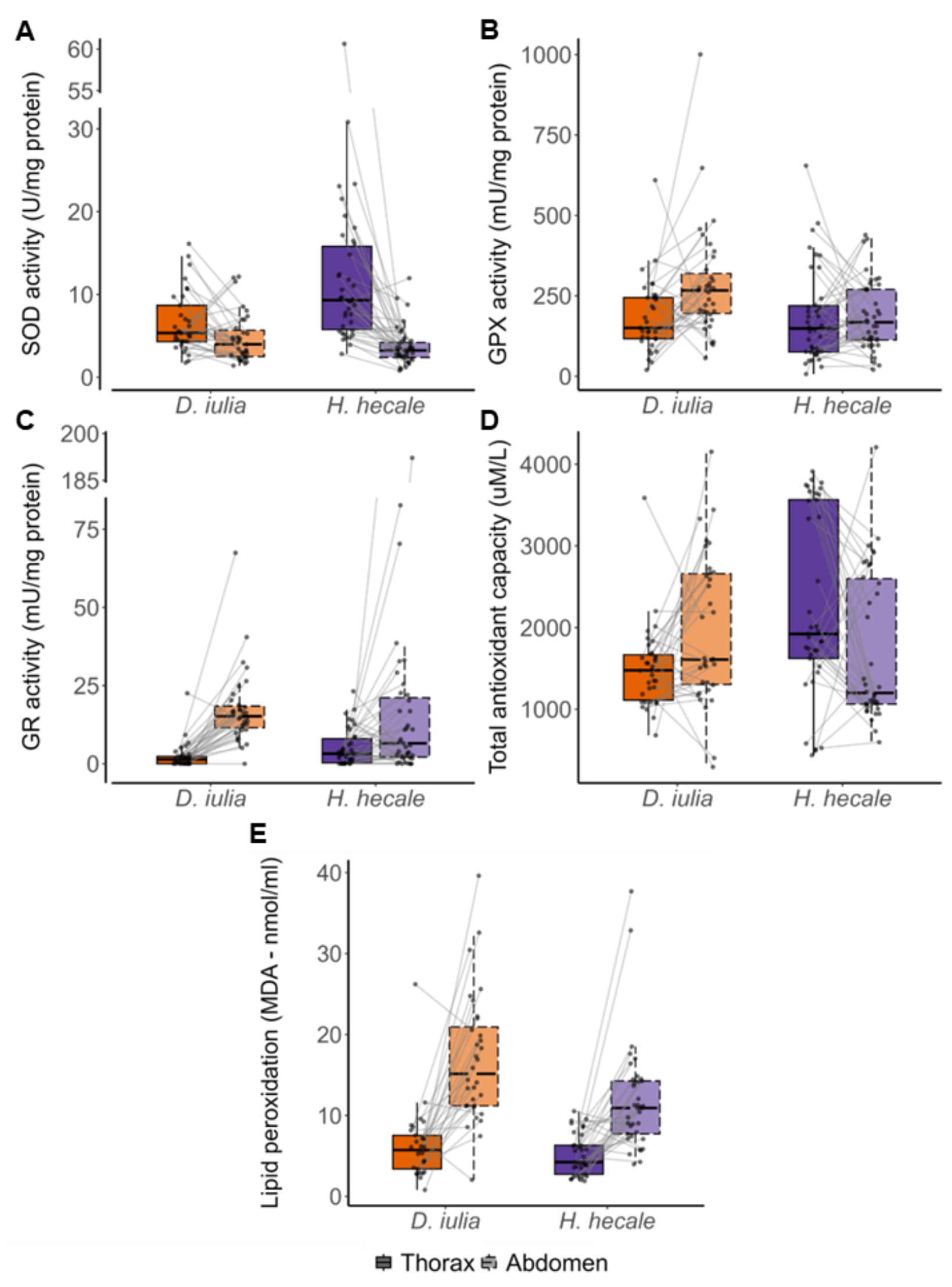
Biomarkers of oxidative stress across tissues in *D. iulia* and *H. hecale*. Measurements of **A** superoxide dismutase (SOD) activity, **B** glutathione peroxidase (GPX) activity, **C** glutathione reductase (GR) activity, **D** total antioxidant capacity (TAC), and **E** lipid peroxidation (malondialdehyde, MDA) in both the thorax (solid outlines, darker fill) and abdomen (dashed outlines, lighter fill) of *D. iulia* (orange) and *H. hecale* (purple). Enzyme activities are normalised to protein content. Points show individual samples, and grey lines connect the thorax and abdomen of the same individual. Boxes show the median and interquartile range, and whiskers extend to 1.5 × the interquartile range. Data are pooled across ages and diets. The y-axes in **A** and **C** are broken to avoid compression due to outliers. *n* = 75 individuals (see Table 2). **Alt text:** Box plots of five oxidative stress markers in thorax and abdomen of both species.

### Thoracic antioxidant capacity associated with tissue preservation in Heliconius

We next sought to investigate whether deviation from expected tissue mass was associated with oxidative stress. We used residuals from the tissue mass model as an index of relative tissue preservation, with positive residuals indicating heavier than expected tissue (relatively more preserved) and negative residuals indicating lighter than expected tissue (relatively more depleted). In *H. hecale*, there was a significant interaction between tissue and residual when modelling TAC (χ^2^_1_ = 13.64, *p* = 0.001), such that a heavier-than-expected mass was associated with increased TAC in the thorax (*t*_70.5_ = 3.73, *p* < 0.001) but not in the abdomen (*t*_70.5_ = −0.08, *p* = 0.935) (Figure 4). This was not the case for *D. iulia*, where TAC was not associated with our index of relative tissue depletion in either tissue (χ^2^_1_ = 0.40, *p*_adj_ = 0.733). Despite this apparent association of antioxidant capacity with tissue preservation in the thorax, relative tissue preservation was also significantly positively associated with levels of lipid peroxidation across both tissues in *H. hecale*, with an even bigger positive association in the thorax (χ^2^_1_ = 6.64, *p*_adj_ = 0.020) (Figure 4), indicating that tissues with higher-than-expected mass tended to exhibit higher levels of lipid peroxidation. None of the other biomarkers measured were significantly associated with our index of relative tissue depletion in either species.

**Figure 4:**
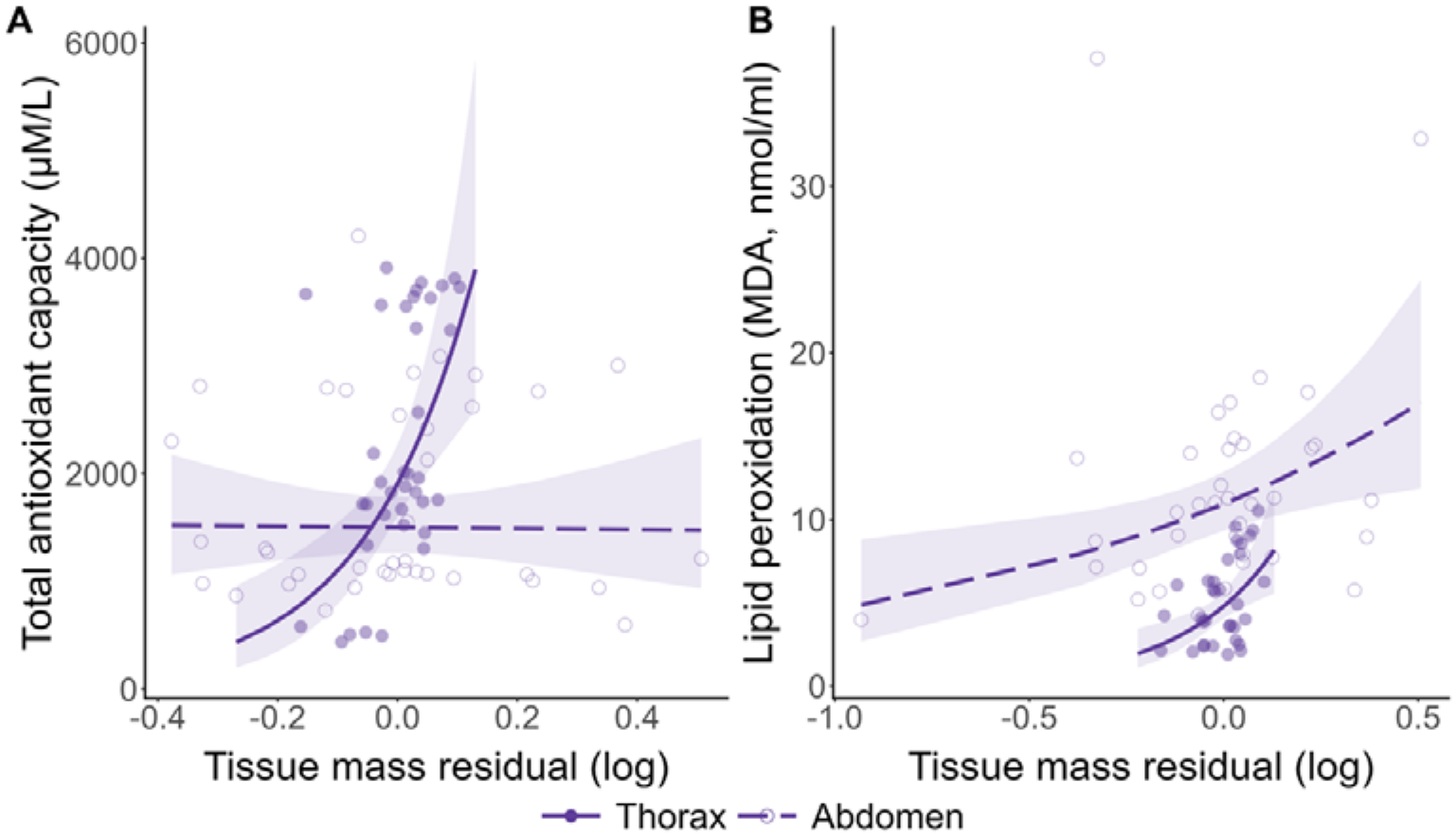
Association between relative tissue preservation and oxidative stress biomarkers in *H. hecale*. Relationship between the tissue mass residual and **A** total antioxidant capacity (TAC) and **B** lipid peroxidation (MDA) in the thorax (filled circles, solid lines) and abdomen (open circles, dashed lines) of *H. hecale*. The residual is the deviation (observed − predicted, log scale) from the mass expected for each tissue given the individual’s age, sex, eclosion mass and freezing status. Positive residuals indicate that a tissue was heavier than expected (less depleted), and negative that it was lighter than expected (more depleted). Points show individual tissue samples, and lines show predictions from the mixed models with 95% confidence intervals, back-transformed to the original scale. *n* = 40 individuals (see Table 2). **Alt text:** Line graphs showing associations of **A** total antioxidant capacity and **B** lipid peroxidation against tissue mass residuals in *H. hecale*. TAC rises with residual in thorax only; MDA rises in both tissues.

## Discussion

The long-lived *Heliconius* butterfly genus is a promising new model system for the comparative biology of ageing, combining experimentally tractable lifespans (Foley *et al*., 2026), a depth of ecological knowledge (Jiggins, 2017), and availability of genomic tools (Concha *et al*., 2019; Cicconardi *et al*., 2023). However, investigations into its lifespan extension have thus far been restricted to ultimate evolutionary mechanisms. Here, we find evidence that enhanced oxidative protection of flight tissue is one candidate proximate mechanism of longevity in this genus.

Our results broadly suggest enhanced oxidative protection of flight tissue in the longer-lived *Heliconius* species. Activity levels of superoxide dismutase, a key antioxidant enzyme, and overall total antioxidant capacity were both higher in the thoraces of *H. hecale* than those of the shorter-lived *D. iulia*, in line with an increased investment in antioxidant defences. Correspondingly, levels of oxidative damage, as measured by lipid peroxidation, were also lower in the *H. hecale* thorax, as well as the abdomen. These results are consistent with the links drawn between longer lifespans and resistance to oxidative stress in a wide range of species (Hulbert, Faulks and Buffenstein, 2006; Salmon *et al*., 2009; Xia and Møller, 2018), with cross-species variation in lifespan thought to be driven particularly by reduction in susceptibility to oxidative damage (Barja, 2002). In its examination of two closely-related species, our study provides a rare insight into the association between oxidative phenotypes and lifespan while minimising the potential confounding effects of broader physiological and ecological differences that may arise across distantly related taxa. Our finding of increased thoracic antioxidant defences in the longer-lived *H. hecale* is also supported by findings from a similar study showing positive associations between TAC and lifespan across closely-related tephritid fly species (Malod *et al*., 2020).

Our analysis of age-related declines in body mass suggested that the observed differences in oxidative phenotypes between *H. hecale* and *D. iulia* are associated with physiological function. In line with the observed thoracic oxidative protection in *H. hecale*, this tissue does not appear to deteriorate with age in this species, in contrast with the age-related decline in *D. iulia* thoracic mass. In fact, relative preservation of thoracic mass is also associated with increased TAC in *H. hecale*, but not *D. iulia*, suggesting a potential mechanistic link – although we cannot infer causality from these associations, as both oxidative stress and tissue mass were measured upon dissection. Interestingly, higher-than-expected tissue mass, particularly in the thorax, was also associated with increased lipid peroxidation in *H. hecale*. One possible explanation is that individuals maintaining greater thoracic mass also retain higher levels of metabolic activity, potentially resulting in both greater antioxidant capacity and increased oxidative damage. However, this interpretation remains speculative, as metabolic activity was not directly measured.

In these butterflies, the thorax is largely composed of muscle tissue, which acts to control movements of the wings and legs. The increased thoracic lipid peroxidation and associated steeper decline in thoracic mass in *D. iulia* observed here aligns with our previous work showing longitudinal age-related declines in muscle function in *D. iulia,* but not in *H. hecale*, as measured by an assay measuring foreleg grip strength (Foley *et al*., 2026). That study also included an investigation of age-related changes in flight behaviour (as measured by time spent flying in a 45-second window), which showed mixed evidence for age-related declines in *D. iulia* but found no evidence for an age-related decline in flight behaviour in *H. hecale* (Foley *et al*., 2026). This is also supported by the lack of an age-related decline in resting metabolic rate in a related *Heliconius* species, *H. charithonia* (Ramos-Pérez *et al*., 2020), at ages up to 45 days old, in contrast with the age-related declines in flight and resting metabolic rate found in several other species in the nymphalid butterfly family, including *Speyeria mormonia* (Niitepõld and Boggs, 2015), *Pararge aegeria* (Vande Velde and Van Dyck, 2013), and *Vanessa cardui* (Woods Jr. *et al*., 2010), (although not in *Melitaea cinxia* (Niitepõld and Hanski, 2013)). Together, these results suggest that the superior protection and maintenance of thoracic muscular tissue demonstrated in *H. hecale* may help to mediate the slowed this slowed physiological ageing and increased lifespan observed across *Heliconius*.

The protection of thoracic muscle tissue in particular in *Heliconius* is significant because of the high metabolic cost of insect flight. Flying insects have resting metabolic rates that are approximately three times as large as those of non-flying insects (Reinhold, 1999), and metabolic rate may increase several hundred-fold during insect flight (Levin *et al*., 2017). Measurements in *Heliconius* specifically show flight and post-flight metabolic rates up to ten times higher than the resting metabolic rate (Salas Prado and Mendoza Cuenca, 2025; Ramos-Pérez *et al*., 2020). In theory, increased metabolic rate may not necessarily be coupled to oxidative damage (Brand, 2000), with one study in cold-exposed voles showing negligible impacts of the experimentally increased metabolic rate on oxidative stress (Selman *et al*., 2008). However, cold exposure in this study resulted in less than two-fold elevations in metabolic rate, a figure that is dwarfed by the elevations induced by insect flight (Levin *et al*., 2017; Salas Prado and Mendoza Cuenca, 2025; Ramos-Pérez *et al*., 2020). Accordingly, several studies across both birds and insects have shown increased oxidative damage associated with active flight (Costantini, Dell’Ariccia and Lipp, 2008; Dick and Guglielmo, 2019; Margotta, Roberts and Elekonich, 2018; Levin *et al*., 2017; Yan and Sohal, 2000; Magwere *et al*., 2006). It seems likely that the accrual of such oxidative damage across a lifetime could potentially limit an insect’s capacity to evolve increases in longevity. In *Heliconius*, the evolution of pollen-feeding may have allowed for greater allocation of larval resources towards antioxidant defences (Foley *et al*., 2026; Dunlap-Pianka, Boggs and Gilbert, 1977), which are thought to be costly (Metcalfe and Alonso-Alvarez, 2010). This may have led to a distinct oxidative phenotype in *Heliconius* characterised by increased protection and maintenance of metabolically-demanding flight tissue, as evidenced by the increased antioxidant activity and reduced oxidative damage in the *H. hecale* thorax, and allowed for the evolution of longer lifespans.

Thorax-specific increases in TAC associated with enhanced survival have also been observed in monarch butterflies after subjection to early-life forced flight (Shephard, Hund and Snell-Rood, 2023). This supports the idea that increased antioxidant defences are potential mechanistic mediators of lifespan extensions in butterflies, although this study did not find any associated differences in oxidative damage in the forced-flight group (Shephard, Hund and Snell-Rood, 2023). The oxidative phenotype in *H. hecale* may also be associated with behavioural adaptations; for example, the wingbeat frequency of *H. hecale* is also about half that of *D. iulia* (Page *et al*., 2024). This might suggest that a lower metabolic rate (Darveau, 2024) may also have contributed to reductions in oxidative damage in *H. hecale*. However, metabolic rate does not always correlate inversely with lifespan in butterflies (Niitepõld and Hanski, 2013), and regardless, the thorax-specific increases in SOD activity and TAC in *H. hecale* suggest that these reductions in oxidative damage are rather related to the superior antioxidant defences of this species. These adaptations may have permitted *Heliconius* to mitigate the accrual of age-related oxidative damage across a lifetime spent flying, allowing it to continue in active flight in late life uninhibited by age-related declines.

In both species, oxidative damage (as measured by levels of lipid peroxidation) was higher in the abdomen than in the thorax, possibly reflecting the metabolic costs of reproduction, which are thought to be high (Metcalfe and Alonso-Alvarez, 2010; Simmons, Lovegrove and Lymbery, 2018). This is supported by the higher GPX activity in the abdomens of *D. iulia* in comparison to *H. hecale*, potentially reflecting a compensatory response to greater oxidative challenge, which may be associated with the higher daily oviposition rate in this species (Dunlap-Pianka, Boggs and Gilbert, 1977; Dunlap-Pianka, 1979). However, TAC is generally thought to be a better readout of oxidative protection than individual antioxidants due to their involvement in various synergistic and antagonistic interactions (Monaghan, Metcalfe and Torres, 2009), and levels of lipid peroxidation are still lower in the abdomens of *H. hecale*, so the link between daily reproductive output and oxidative stress is not straightforward. In any case, some researchers have cautioned against over-interpretation of within-individual differences in oxidative stress between tissues, citing the differences between tissues in types and levels of ROS (Selman *et al*., 2012), antioxidant machinery, and capacity and mechanisms of repair (Halliwell and Gutteridge, 2015), as well as an observed lag in response to oxidative stress between tissues (Costantini, 2019). The observed differences in lipid peroxidation between thoraces and abdomens of both species may therefore be a poor proxy for the costs of specific physiological functions.

The lack of an effect of age on oxidative markers in either butterfly was unexpected, considering the evidence for age-related increases in oxidative stress in many species, including in other flying insects (Savić *et al*., 2026; Magwere *et al*., 2006). This may simply be because we were underpowered, with a relatively small sample size (*N_H. hecale_* = 40, *N_D. iulia_* = 35), or possibly due to selective disappearance of more oxidatively-stressed individuals from older age classes, an unfortunate limitation of our necessarily cross-sectional approach (Speakman *et al*., 2015; Selman *et al*., 2012). However, the effects of age on markers of oxidative stress are not always consistent. One study found evidence for increased oxidative damage with age in only one of the four social insect species studied, and only in the worker caste (Kramer *et al*., 2021). Likewise, across tephritid fly species no effect of age on markers of oxidative damage or antioxidant protection was found, despite finding differences in TAC correlating with lifespan (Malod *et al*., 2020). We argue that here the absence of robust age-related changes in oxidative markers suggests that the observed interspecific differences represent relatively stable physiological differences between *H. hecale* and *D. iulia* rather than divergent patterns of oxidative ageing.

We also found no evidence for an effect of diet on oxidative markers, which was unexpected considering the evidence for relationships between diet and oxidative status across species (Metcalfe and Alonso-Alvarez, 2010; Haddad, Kelbert and Hulbert, 2007). Our previous work showed a diet-dependent lifespan extension in pollen-fed *H. hecale*, which we thought could be mediated via contributions of pollen-derived nutrients to oxidative stress resistance. The lack of a diet effect may once again be due to insufficient power. However, it may also simply be the case that access to dietary pollen during adulthood has no effect on oxidative markers in *H. hecale*, and the observed inter-specific differences in oxidative phenotypes are mediated solely through differences in larval resource allocation, as suggested above. The absence of an effect of adult diet, or indeed age, on oxidative markers in either butterfly supports this, suggesting that these oxidative phenotypes are fixed at some point during development, and remain stable throughout adulthood. Future work on oxidative stress in the larval and pupal stages could help to assess whether these phenotypes differ across life stages, as has been observed in amphibians (Martin *et al*., 2024).

## Conclusions

A particular strength of the *Heliconius* system is the framework offered by comparisons with their shorter-lived Heliconiini close relatives, from whom they diverged relatively recently in evolutionary time (∼18 mya) (Cicconardi *et al*., 2023). The ecological, physiological, and phylogenetic similarities between butterflies in the Heliconiini tribe allows for the association of differences in oxidative physiology to their pollen-feeding behaviour and associated lifespan differences, rather than other derived traits. Future analyses including a larger spread of species from across the Heliconiini tribe would help to test the consistency and validity of these associations. These associations would also benefit from experimental validation, for example by testing resistance to exogenous stressors such as paraquat, a herbicide which acts to induce acute oxidative stress (Arking *et al*., 1991), or similarly 2,4-dinitrophenol or diclofenac. Measurement of oxidative markers in wild-caught individuals would also help to assess the evolutionary relevance of these differences in nature (Selman *et al*., 2012), where butterflies fly across much larger distances (Mallet, 1986) in search of food, mates, and oviposition sites. While these results offer the first promising plausible pro-longevity mechanism in *Heliconius*, ageing is a multifarious phenomenon, by its nature implicating whole-organism systems and processes. We therefore expect and welcome the future studies likely to identify complementary, and perhaps intertwined, mechanisms of longevity in *Heliconius*.

## Acknowledgements

We would like to thank the Ministerio del Ambiente in Panama for permission to collect butterflies used in these experiments. We would also like to thank the Smithsonian Tropical Research Institute, and in particular Owen McMillan, for providing the facilities to carry out much of this research. We are grateful for the efforts of Cruz Batista, Greta Hernández, Josie McPherson, and Made Roger in rearing butterflies at the insectaries, as well as to Francisco Miranda from the Ecophysiology Lab of EBD-CSIC for his assistance with biochemical analyses. Generative AI was used to assist with writing code, primarily for plotting purposes. This work was supported by a DTP Studentship as part of the GW4 BioMed MRC DTP (MR/N0137941/1) awarded to the Universities of Bath, Bristol, Cardiff and Exeter from the MRC/UKRI as well as a Travelling Fellowship from The Company of Biologists (JEBTF23101308) (J.F.). It was also supported by a NERC Independent Research Fellowship (NE/N014936/1) and ERC Starter Grant (758508) (S.H.M.), as well as a Ramón y Cajal fellowship (2023-044964-I) (Spanish Ministry of Science, Innovation, and Universities) (P.B.).

