## Supplementary Material for "An elongated lifespan is associated with higher oxidative protection of flight tissue in Heliconiini butterflies"

Supplementary Figures

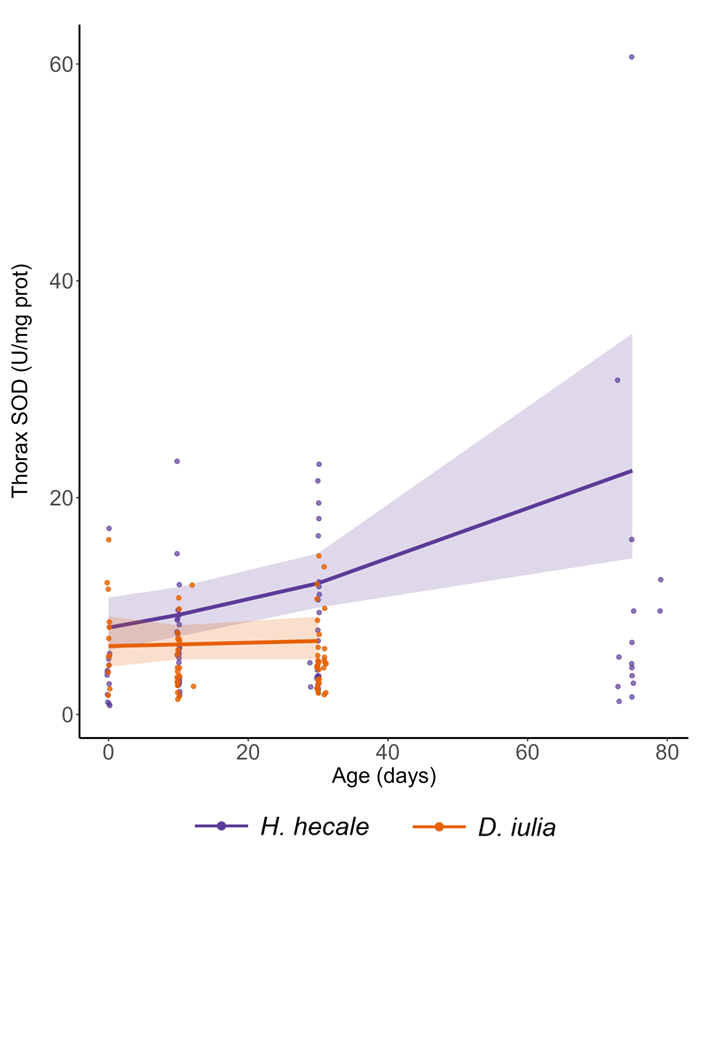

**Figure S1: Thorax SOD activity with age.** Superoxide dismutase (SOD) activity (U/mg protein) in the thorax of *H. hecale* and *D. iulia* as a function of age. Points show individual samples; note the large outlier in the oldest *H. hecale* age group, which biases the model predictions (see Results). Lines show predictions from the generalised linear models for each species, with shaded 95% confidence intervals. Data are pooled across diets and sexes. **Alt text:** Line graph of thorax SOD activity plotted against age for both *H. hecale* and *D. iulia*.

Supplementary Notes

Supplementary Note 1

As mentioned in the Materials and Methods section, body and tissue-specific mass data was collected for 85 individuals of *H. hecale* and 62 individuals of *D. iulia*. However, tissues from only 40 individuals of *H. hecale* and 35 individuals of *D. iulia* were then snap-frozen, stored in a −80° freezer, and transported to the U.K. and then Spain for oxidative stress analyses. For this oxidative stress group, individual tissues (thorax/abdomen) were weighed just before homogenisation, after retrieval from the freezer. For the remaining individuals, which were not snap-frozen and transported elsewhere but rather retained for other analyses, individual tissues were weighed fresh, immediately following decapitation. The freeze-thaw cycle appears to have affected weights of the abdomens but not the thoraces, with “Frozen” abdomens weighing less than “Fresh” abdomens (Figure S1), likely due to this tissue’s higher liquid content, which may have been partially lost upon defrosting. This result was found in both *H. hecale* and *D. iulia*. A tissue:frozen interaction was included in all tissue-specific mass analyses to account for this.

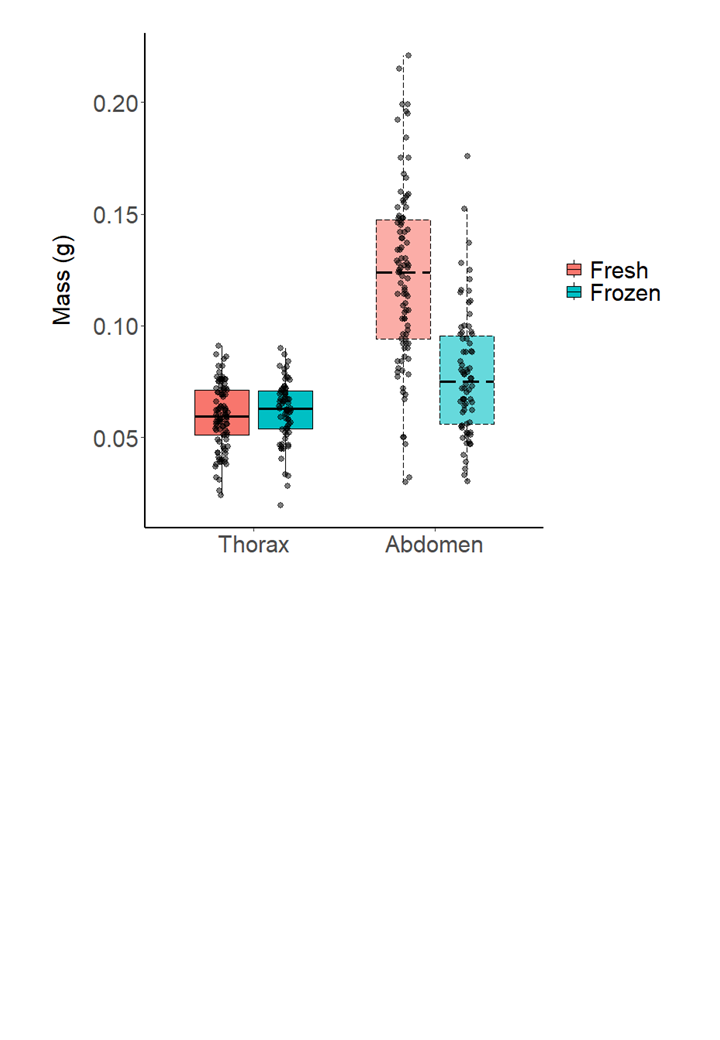

**Figure S2: A single freeze-thaw cycle is associated with lower recorded values for mass in the abdomen specifically.** *n* = 175 individuals (see Table 1), each with values for both the thorax and the abdomen. **Alt text**: Boxplots of tissue-specific mass, differentiating between samples that underwent a freeze-thaw cycle before recording weight and samples which were weighed fresh.
